# Elucidating key determinants of engineered scFv antibody in MMP-9 binding using high throughput screening and machine learning

**DOI:** 10.1101/2024.06.04.597476

**Authors:** Masoud Kalantar, Ifthichar Kalanther, Sachin Kumar, Elham Khorasani Buxton, Maryam Raeeszadeh-Sarmazdeh

## Abstract

An imbalance in matrix metalloproteinase-9 (MMP-9) regulation can lead to numerous diseases, including neurological disorders, cancer, and pre-term labor. Engineering single-chain antibody fragments (scFvs) Targeting MMP-9 to develop novel therapeutics for such diseases is desirable. We screened a synthetic scFv antibody library displayed on the yeast surface for binding improvement to MMP-9 using FACS (fluorescent-activated cell sorting). The scFv antibody clones isolated after FACS showed improvement in binding to MMP-9 compared to the endogenous inhibitor. To understand molecular determinants of binding between engineered scFv antibody variants and MMP-9, next-generation DNA sequencing, and computational protein structure analysis were used. Additionally, a deep-learning language model was trained on the synthetic library to predict the binding of scFv variants using their CDR-H3 sequences.

## Introduction

Matrix metalloproteinases (MMPs) are a family of proteinases with central roles in remodeling and regulating the extracellular matrix ^1,2^. MMPs are associated with several diseases when not regulated by their inhibitors and activators ^1,2^, making them suitable targets for developing novel protein-based therapeutics ^1,3^. The catalytic domains of MMP family members share high amino acid similarity and their active sites are extensively conserved. Consequently, developing small molecule inhibitors to distinguish different MMPs is extraordinarily difficult and was not successful in progressing to the late stages of clinical trials due to their low stability and low binding selectivity, which caused toxicity and side effects ^3^. Thus, the attention to developing therapeutics targeting MMPs has been recently changed to protein alternatives.

Monoclonal antibodies (mAbs) are well-established protein therapeutics generally well-tolerated by the body with minimal risk of side effects. ^4^. Various antibodies have been previously used ^5–7^ and engineered ^8–10^ to target proteases, especially MMPs. mAbs provide a large protein-protein (antigen-antibody) binding interface area along with multiple flexible binding loops and complementarity-determining regions (CDRs) ^11^. Antibody-based therapeutics also offer high binding affinity and selectivity, which can be further enhanced using protein engineering and design strategies, such as directed evolution and yeast surface display ^12–14^. Moreover, antibody production and purification platforms are well-established, and the antibody-based drug industry and market are already highly developed ^12,15,16^. These reasons make antibodies an excellent protein scaffold for engineering MMP inhibitors. Among the MMPs, the overexpression of MMP-9 has been strongly linked to a poor prognosis in various malignancies, including breast, lung, colon, gastric, pancreatic, and prostate cancer ^17,18^. Further, advances in generating humanized antibodies, and various antibody fragments, such as antigen-binding fragments (Fab), single chain antibody fragments (scFv), and camelid antibodies (CAb), as recombinant proteins ^19^ makes antibodies one of the leading classes of biological binding molecules, easily cloned, produced, and genetically manipulated for diverse purposes. Antibody discovery efforts to find potential therapeutics targeting MMPs, specifically MMP-9 due to its pathological contribution in several diseases, led to several hit antibodies ^20,21^.

One of the first antibodies developed to target MMP-9 was a murine mAb, REGA-3G12 ^20^. REGA-3G12 interacts with the N-terminal region of the catalytic domain however it does not interact directly with the zinc-binding site ^22,23^. The Complementarity-Determining Region 3 of the Heavy Chain (CDR-H3) region has been identified as a key antigen recognition site and a crucial loop in enhancing antigen binding strength ^24,25^. However, the probability of producing antibodies from humans or murine that target MMPs effectively is difficult because active sites of MMPs are often deeply buried within a major cleft or concave structure, making it inaccessible to the antigen-binding sites of native human antibodies. These antibodies typically feature a flat surface, with the CDR-H3 averaging 9 to 12 amino acids in length. To overcome these limitations, several synthetic human antibody libraries were constructed by strategically incorporating antigen-binding regions (paratopes) derived from camelid antibodies, as they have long CDR-H3 regions and can penetrate the concave shape of the enzyme active site ^26^. These results suggest that at least for active site inhibitors, the appropriate distribution of CDR-H3 lengths and amino acid compositions could be a significant factor in accommodating desired paratope conformations that are compatible with the topology of the targeted MMPs.

The advancement of machine learning (ML), particularly deep learning (DL) and natural language processing (NLP) technologies, along with increased computing power, has further enhanced biotechnological applications, including protein design and engineering ^27–31^. These developments have led to the creation of Large Protein Language Models (LPLMs), which assist in discovering the evolutionary, structural, and functional properties across protein space by encoding amino-acid sequences into numeric vector representations ^32^. In this study, we leveraged pretrained LPLMs to extract features from CDR-H3 sequencing data to train a downstream Long-Short-Term Memory (LSTM) model to predict the binding affinity between CDR-H3 and MMP-9cd. To understand the predictive influence of each amino acid in CDR-H3 on the binding affinity, we used SHapley Additive exPlanations (SHAP). This technique applies game theory to explain the contribution of each input feature to the prediction made by an ML model. By analyzing the file generated by this technique, we can gain insights into feature importance, understand the distribution of feature impacts, detect interactions between features, and interpret the overall behavior of the model.

This study refines a synthetic scFv antibody library previously engineered for decreasing non-specific binding ^24^ to improve binding to the MMP-9cd (**Fig. 1**). We achieved this by combining the fluorescent-activated cell sorting (FACS) screening approach with next-generation sequencing (NGS) analysis. FACS allows for efficient identification of high-affinity binders, while NGS offers an unparalleled level of detail compared to traditional methods. By providing both the sequence and frequency information for each scFv antibody in the library, NGS allows us to pinpoint key amino acid residues crucial for binding to MMP-9cd. The results of negative (non-binders) and positive (binders) using FACS sorts were used to fine-tune LPLMs. These ML-developed models focused on CDR-H3 could predict not only the scFv variants kept as testing population with high precision, but also the other negative and positive MMP-9 binders that were unrelated to this study or library. This comprehensive approach goes beyond traditional screening techniques, enabling the identification of promising candidates that might otherwise be missed.

**Figure 1.**
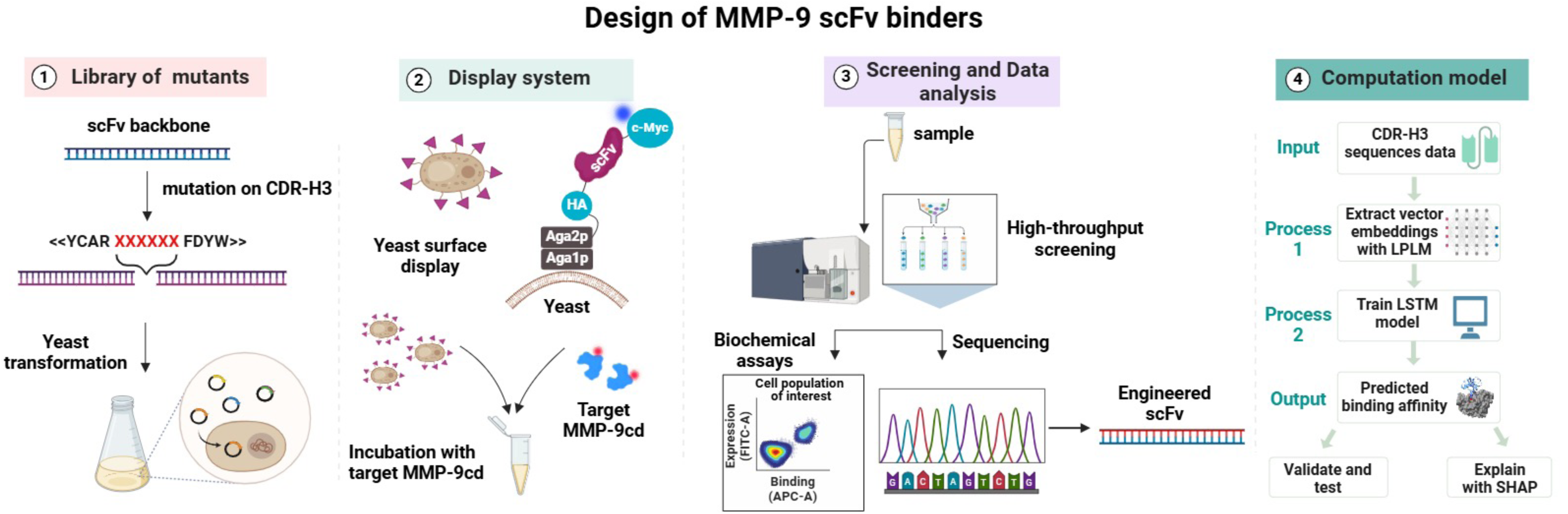
The general approach for protein engineering and design of antibody scFv variants using directed evolution, yeast surface display to target MMP-9. 1) Library generation: a library of scFv variants is created by heavily mutating the CDR-H3 region to introduce diversity in both amino acid composition and length. These variants are then electrotransformed into a robust yeast strain for further selection. 2) Expression and display: yeast cells carrying expression plasmid vectors encoding different scFvs can be cultured and induced to secrete these scFvs. A signal peptide within the vector directs the scFvs to the secretory pathway, where Aga2p forms a disulfide bond with Aga1p on the yeast cell wall, facilitating the display of each scFv variant on the cell surface. Cells expressing different scFvs are then incubated with the target MMP-9 enzyme for further analysis. 3) Screening and sequencing: the scFv library is screened and sorted for binders and non-binders using FACS. The sorted libraries or isolated scFvs are then sequenced via next-generation sequencing and Sanger sequencing. 4) Data analysis and model training: sequencing data extracted from NGS is used as input for a machine learning model analyzing the effect of CDR-H3 on MMP-9 binding. This data is used to train and validate a model that can predict the binding affinity of specific CDR-H3 regions to MMP-9.

## MATERIALS AND METHODS

### Strain and plasmids

*Saccharomyces cerevisiae*, strain EBY100 (aGAL1-AGA1:: URA3 ura3 52trp1leu2Δ200pep4::HIS2prb1Δ1.6Rcan1 GAL) and RJY100 strain ^24^ were used for yeast surface display of the N-terminal domain of TIMP-1 (N-TIMP-1) as the control, and scFv variants in the naïve library, respectively. The N-TIMP-1 is expressed at the N-terminus of the Aga2p protein and subsequently integrated into the pCHA vector backbone, resulting in a free N-terminus capable of binding to MMP-9cd. As for scFv variants, the pCTCON2 vector was employed to express the scFv at the C-terminus of Aga2p ^24^.

### Yeast surface display of N-TIMP-1 and scFvs

The yeast cells, which had been electroporated with plasmids, were incubated overnight in minimal SDCA, pH 6. This media composition included 20 g/L dextrose, 6.7 g/L yeast nitrogen base, 5 g/L Bacto casamino acids, 10.19 g/L sodium phosphate dibasic (Na_2_HPO_4_·7H_2_O), and 8.56 g/L sodium dihydrogen phosphate monohydrate (NaH_2_PO_4_·H_2_O) and in a shaker incubator at 30°C and 250 rpm. Subsequently, the yeast cells were induced in SGCAA media (similar to SDCAA but with 20 g/L galactose instead of dextrose, pH 6) for 20 h at 30°C, starting with an initial OD600 of 1.00. Yeast cells displaying N-TIMP-1 or scFv variants were harvested at an OD600 of 0.2 and washed twice in 750 µL of ice-cold PBSA (8 g/L NaCl, 0.2 g/L KCl, 1.44 g/L Na_2_HPO_4_, 0.24 g/L KH_2_PO_4_, pH 7.4, and 0.1% BSA). Subsequently, these cells were incubated with His6x-MMP-9cd from Enzo Life Sciences (Farmingdale, NY), a final concentration of 400nM, for 60 minutes at room temperature, following previously reported methods ^33^. After incubation, the samples underwent two washes with 750 µL of ice-cold PBSA, and the cells were kept on ice. For the primary antibody labeling process, mouse anti-c-Myc Antibody (GenScript, NJ) solution at a 1:100 ratio in PBSA buffer (0.25 mg/ml) was used to incubate the cell for 30 minutes on ice. Subsequently, for the secondary antibody labeling, the cells washed three times in ice-cold PBSA were incubated with goat anti-mouse Alexa Fluor 488 antibody (Invitrogen, 2mg/mL) and anti-His6x monoclonal antibody conjugated with Alexa Fluor 647 (Invitrogen, 1mg/mL) at 1:100 dilution in PBSA for either one for 30 minutes on ice, shielded from light. Following the completion of the final washes, the labeled cells were resuspended in 750 µL of PBSA for analysis using a BD Accuri^TM^ C6 Plus flow cytometer (BD Biosciences, NJ). The data obtained from the flow cytometer were further analyzed using FlowJo software (FlowJo, LLC, OR).

### Protein expression and purification

Recombinant active human MMP-9cd (auto-cleavage resistance) ^34^ with an N-terminal His6x tag was expressed in Rosetta™(DE3) pLysS (Millipore Sigma) cells using a pET-28a(+)-His6x-MMP-9cd vector as previously described ^33,35,36^. Briefly, MMP-9cd protein expression was induced with 0.5 mM IPTG for 21 hours at 25°C. Insoluble MMP-9cd protein was extracted from E. coli cells via sonication and solubilized under denaturing conditions with urea. Bacterial pellets were mixed with lysis buffer and kept shaking overnight at 4°C. Lysate was then incubated with sodium deoxycholate and DNase I for 1 hour at room temperature. The lysate was then separated by centrifugation, and aliquots were collected for subsequent SDS-PAGE analysis. This cycle of sonication, solubilization, lysis, and centrifugation was repeated until no insoluble material remained in the pellet fraction. The pooled supernatants containing solubilized MMP-9cd were purified by immobilized metal affinity chromatography (IMAC) using Ni resin. Finally, the purified MMP-9cd was refolded through gradient dialysis. For the original sorts, purchased His6x-MMP-9cd protein from Enzo Life Sciences (Farmingdale, NY) was used.

### Screening the yeast-displayed scFv library using FACS

scFv library, a generous gift from the Wittrup lab-MIT-Chemical Engineering, was previously engineered to minimize the non-specific binding to various protein targets ^24^. The library’s structure consisted of 5 different heavy chain segments (V_H_) and 3 variations of light chains (V_L_), connected by a glycine linker, all presented in the V_L_-V_H_ format. To expand the diversity of the library, heavy mutations were applied to the CDR-H3 loop in terms of length and amino acid compositions ^24^.

The library of synthetic scFv library was originally recovered from the glycerol stock in 50 ml SD-CAA media (20 g glucose, 6.7 g yeast nitrogen base without amino acids, 5 g casamino acids, 10.4 g sodium citrate, 7.4 g citric acid monohydrate, pH 4.5) containing 100 µg/mL ampicillin to prevent bacterial growth. The library was diluted to 500-1000 L after overnight growth at 30°C shaker. Before each round of cell sorting, the yeast cells were induced in SGCAA media. The number of cells was measured to reach a density of OD_600_ of 1.00 (10^^7^ cells/ml). Subsequently, the cells were incubated with His6x-MMP-9cd, followed by primary and secondary antibody labeling. After washing, the cells were resuspended in ice-cold PBSA buffer, following the procedure outlined in the “Yeast surface display of N-TIMP-1 and scFvs” section. These samples were kept on ice and protected from light until they were loaded onto the BD FACS Aria II cell sorter. A pentagon gate was applied to isolate clones that exhibited strong signals for Alexa Fluor-488 and Alexa Fluor-647, indicating expression and positive binding to MMP-9cd, respectively. Additionally, another rectangular gate was used to collect clones with strong signals for Alexa Fluor 488 and weak signals for Alexa Fluor 647, indicating non-binders to MMP-9cd.

After sorting, the cells were recovered by inoculating them in a 50 mL SDCAA medium with a pH of 4.5. They were then allowed to grow overnight at 30°C. The yeast library was stored in 20% glycerol stock at −80°C for longer storage. The library underwent two rounds of screening, each involving staining, sorting, recovery, and regrowth. In the first round, cells were incubated with 300nM MMP-9cd and sorted based on yield mode to eliminate unwanted variants. Approximately 5% of the population (10^^6^ cells) was collected for the positive gate (binders), while up to 3% (3×10^^5^ cells) were collected for the negative gate (weak or non-binders). In the second round, the concentration of MMP-9cd was reduced to 100nM, and the sort mode was changed to purity to increase the efficiency of sorted cells, which exhibit both positive expression and binding signals. Up to 2% of the population (5×10^^5^ cells) was collected in the last sort.

### DNA isolation for Sanger and Next-generation sequencing

After sorting using FACS, isolated yeast clones were grown on selective SDCAA plates at a 30°C incubator for further binding analysis and DNA sequencing. For the Sanger sequencing of isolated clones, the yeast DNA plasmids were extracted using the Zymoprep Miniprep II kit (Zymoprep), amplified using PCR, and purified using the SV Gel and PCR Clean-Up System (Promega Corporation) and were sequenced at the Eurofins genomics.

For the next-generation sequencing (NGS) analysis, the yeast plasmid DNA from the scFv antibody libraries, either negative or positive sorts, were isolated using the Zymoprep Miniprep II kit. To ensure high-quality sequencing data, the plasmid DNA underwent Lambda-Exo digestion to remove impurities, including yeast cell genomes, from the extracted DNA as previously described ^37^. Subsequently, the heavy chain genes were selectively amplified by PCR using Phusion high-fidelity polymerase and corresponding primers that were up and downstream of CDR-H3. This targeted amplification strategy was chosen due to the high mutation rate in the CDR-H3 and further analysis of this region. Finally, the amplified scFv variants (positive or negative) were sequenced by Azenta/Genewiz using an Illumina sequencing platform.

DNA sequences obtained from NGS were analyzed using multiple Python scripts. These scripts were designed to sort each V_H_ variation and align the CDR-H3 regions. The Python codes included the translation of DNA sequences into three forward and three reverse frames of amino acid sequences, the identification of amino acid residue mutations in the CDR-H3 region, and counting the number of repetitions. Subsequently, the frequency of amino acid residues in the CDR-H3 region and the CDR-H3 length were compiled and graphed using PRISM software packages (GraphPad Software, Inc., CA).

### Training a DL model for predicting binding

DL models were trained using 2380 unique CDR-H3 sequences of scFv variants in the positive gate and 153 unique CDR-H3 sequences in the negative gate. Two large protein language models (LPLM) were used for extracting features from protein sequences: The Evolutionary Scale Modeling (ESM)-2 models (with 650B, 3B, and 15B parameters) ^27^ and the AntiBERTy model with 26M parameters ^38^. ESM-2, a state-of-the-art LPLM developed by Meta AI, is trained on sequences from the UniRef protein sequence database using a Masked Language Modeling (MLM) objective. In MLM, amino acids are randomly masked in a protein sequence, and the model is trained to predict the missing amino acids from their surrounding context. This approach helps the model learn vector representations (called embeddings) that capture the patterns, dependencies, and structures of protein sequences. The accuracy of structure prediction improves with the model’s size (number of parameters) and the diversity of its training database ^27^. AntiBERTy is similar to the ESM-2 model but has fewer parameters and was trained exclusively on antibody sequences, capturing the diversity and specificity of the immune repertoire.

A low-dimensional visualization of features was extracted from each LPLM model. The embeddings produced by the LPLM for each amino acid in a sequence were averaged over CDR-H3 positions (**Fig. S1A**). Then, the principal components were extracted from the embedding vectors and used for two-dimensional projection using the t-SNE algorithm ^39^. The non-binders (orange dots) form a cluster in the embedding space, demonstrating the utility of LPLMs for distinguishing between CDR-H3s that are binders and non-binders (**Fig. S1B**). Embeddings extracted for each amino acid position in CDR-H3 were padded to the same length and used to train a downstream LSTM model to predict binding affinity with MMP-9cd.

Embeddings extracted for each amino acid position in CDR-H3 were padded to the same length and used to train a downstream LSTM model to predict the binding affinity between CDR-H3 and MMP-9cd. The sequences in the dataset were split into 80% training and 20% test sets. Each model was trained for 50 epochs with early stopping and the Adam optimizer. The initial learning rate was 0.001, with decay steps of 1000 and a decay rate of 0.9. The predictive performance of the model was evaluated using precision, recall, and F1 metrics in both a cross-validation setting and an independent out-of-sample test set.

To understand the predictive influence of each amino acid in CDR-H3 on binding affinity, SHapley Additive exPlanations (SHAP) were employed. SHAP, a technique based on game theory, explains the contribution of each input feature to the prediction made by a machine learning (ML) model ^40,41^. The DeepSHAP algorithm ^40^ was employed to compute the predictive importance of each amino acid residue in CDR-H3 binding. DeepSHAP uses reference baselines to compute differences in neuron activations and linearly decomposes the model’s output into contributions from each input feature. These contributions are then aggregated over multiple reference baselines to capture the effect of all possible feature combinations, providing a scalable and accurate approximation of Shapley values for deep learning models.

Contributions were averaged over the embedding dimension to produce one Shapley value per amino acid residue in a CDR-H3, and SHAP force plots were then created for each sequence. The force plots visualize the impact of each residue on the binding of a single CDR-H3, showing how the residue pushed the prediction for CDR-H3 from the base value to the model output. To understand the overall importance of each position or residue in binding, global Shapley values were computed by averaging the Shapley values over all CDR-H3 samples in the dataset. This average value serves as a clear indicator of how critical a particular position in the CDR-H3 region is for binding. Positions with high average Shapley values are particularly important for binding affinity. Conversely, positions with low average Shapley values have minimal or less consistent impacts on binding affinity. These positions likely do not contribute significantly to the binding process across different possible residues.

The analysis of residue-specific impact reveals how each amino acid residue, averaged across the entire CDR-H3 region, affects binding affinity. This approach identifies residues that generally have positive or negative effects on binding, regardless of their specific location within the sequence. Positional variability refers to the extent of fluctuation in Shapley values for each position within the CDR-H3 region. High variability at a position indicates that its contribution to binding affinity can significantly differ depending on the amino acid present. Such positions are likely crucial for binding interactions with MMP-9cd and should be targeted for optimization.

### AlphaFold2 protein complex modeling and analysis

AlphaFold2-Multimer pipeline was utilized to predict the complete three-dimensional structure of the MMP-9cd-scFv complex ^42^. This approach involved incorporating the amino acid sequences of both the antibody heavy and light chains alongside the MMP-9cd sequence within a single FASTA file. Subsequently, the generated structures were subjected to a relaxation process using Amber, a molecular dynamics package, to relieve steric clashes and optimize the geometry for better physical realism. Also, up to 20 template hits were allowed during the modeling process to enhance the prediction accuracy. These templates provide structural references that guide the folding and interaction predictions.

## RESULTS

### Screening of the synthetic scFv library for binding to MMP-9cd showed improvement in both expression and binding levels compared to the naïve library

The previous analysis identified an enrichment of four amino acids (Gly, Val, Trp, and Arg) within the CDR-H3 region ^24^. Notably, the effect of Trp in CDR-H3 was most dominant as a driver of nonspecificity. This knowledge was subsequently applied to generate a new library (also known as naïve library in this study), resulting in scFv antibodies exhibiting robust binding to a wide range of antigens with minimal nonspecificity. The overall structure of the library uses five V_H_ and three V_L_ frameworks, with the majority of diversity focused on the CDR-H3 loop. Key highlights of this scFv library included the elimination of Trp and a significant drop in the frequency of Arg and Val. Additionally, allowing CDR-H3 loop length diversity (6 to 17 aa), which mimics the natural repertoire, and ensuring a library size of at least one billion members were among the significant improvements ^24^. The synthetic scFv antibody library was subjected to two rounds of FACS targeting MMP-9cd. Two sort gates (positive and negative) were used to collect the cells sorted as binders with dual positive expression and binding signals, and negative binders with only positive expression determined by c-Myc detection and low MMP-9cd binding (**Fig. 2A**, **Fig. S2**).

**Figure 2.**
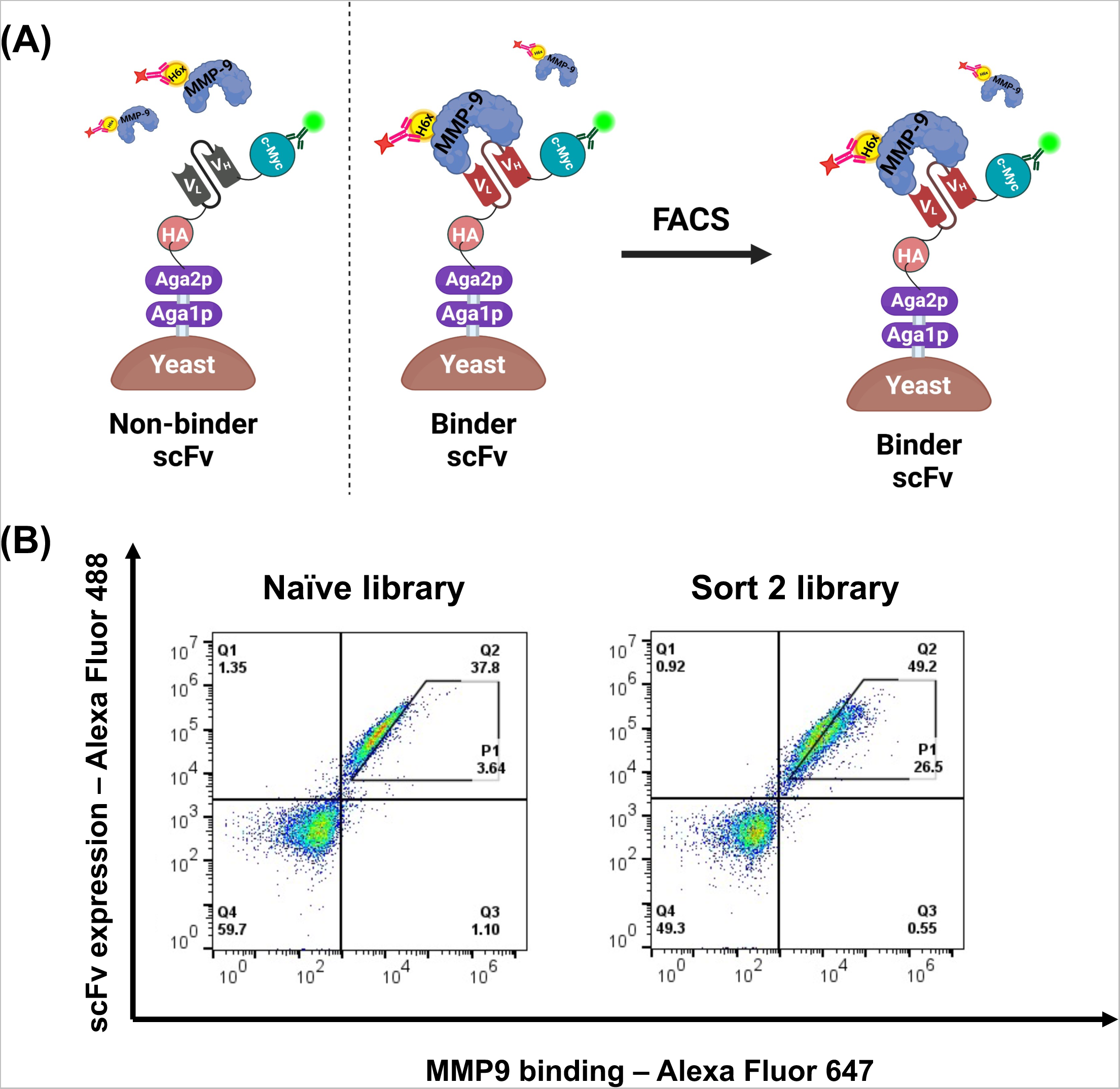
FACS sorting of yeast cells displaying scFvs. A) Yeast cells displaying scFvs are directly incubated with fluorescently labeled MMP-9, enabling efficient sorting and isolation of high-affinity binders using FACS. This process allows for the identification and selection of scFvs with strong binding affinity while discarding those with weak or no binding. B) Flow cytometry analysis reveals a clear difference in MMP-9cd binding between the naive scFv library and the scFv library after two rounds of FACS sorting. The naive library shows a weaker signal, while the sorted library exhibits a stronger signal, demonstrating the effectiveness of the selection process. This is evidenced by the increased fluorescence intensity of MMP-9cd binding (x-axis) in the sorted library. The diagonal gate (P1) defines the enriched population of yeast cells displaying scFvs with high MMP-9cd binding affinity.

Although the scFv antibody framework includes a combination of different light chains and heavy chains, the backbone and other CDR regions except for CDR-H3, which was heavily mutated, exhibit significant similarity to each other with up to 80% sequence homology across the entire scFv backbone. The rationale behind this approach is based on previous studies demonstrating that CDR-H3 serves as the primary functional contributor to antigen recognition in most antigen-binding sites ^43–45^. Additionally, separate selections were conducted to isolate high-affinity binders using the positive gate and to identify weak or non-binders through the negative gate. The number of selection rounds was intentionally limited, aiming not only to isolate the few tightest binders but also to obtain a diverse set of unique binders for comprehensive statistical analysis.

A significant enrichment of positive scFv binders was observed compared to the original or naive library. An increase of more than 10% in the overall expression of collected cells through the positive gate was noted, along with a similar increase in binding affinity (**Fig. S3**). The percentage of positive binders collected after two rounds of sorting in the P1 gate increased significantly from 3% to 26%. This enrichment suggests that the sorting strategy effectively isolated high-affinity binders from the naïve scFv library (**Fig. 2B**).

### Next-generation sequencing of the screened scFv antibody library revealed enrichment of mainly polar residues after two rounds of FACS

Sequence analysis of scFv binder and non-binder variants obtained from positive and negative sort gates showed amino acids that are most likely favorable to contribute to high binding affinity towards MMP-9cd. The NGS data analysis from 2380 unique CDR-H3 sequences in the positive gate and 153 unique CDR-H3 sequences in the negative gate showed an increase in the amino acid frequency in the CDR-H3 sequences of the final positive library compared to the naive library, with a notable increase in Arg frequency by approximately 6-fold (**Fig. 3A**, **Fig. S4**). This observation is particularly interesting considering that the naive library was designed with a low frequency of this residue. The significant enrichment of Arg within the binder-selected pool suggests that its presence in the CDR-H3 loop may aid the folding or structural stability of the heavy chain. This interpretation is supported by the positive correlation observed between the expression level of antibodies on yeast cells and their stability ^46^ since all collected scFv variants in the positive gate exhibit high expression levels. However, the enrichment of positively charged Arg residues could be also attributed to binding to the negatively charged MMP-9cd surface, particularly to the enzyme’s active site and neighboring regions. These findings are consistent with previous studies on CDR-H3 of high-affinity antibody binders, which have shown that Arg residues increase after affinity maturation, and a high Arg content in the CDR-H3 correlates with increased nonspecific binding to the target antigen. ^47^. Even though Arg side chains undoubtedly contribute favorably to binding energy in many protein-protein interactions ^48^. In addition to Arg, other residues that were enriched in the positive pool are mainly polar residues or those with polar side chains, such as Ser, Tyr, Gly, and Gln (**Fig. 3B**).

**Figure 3.**
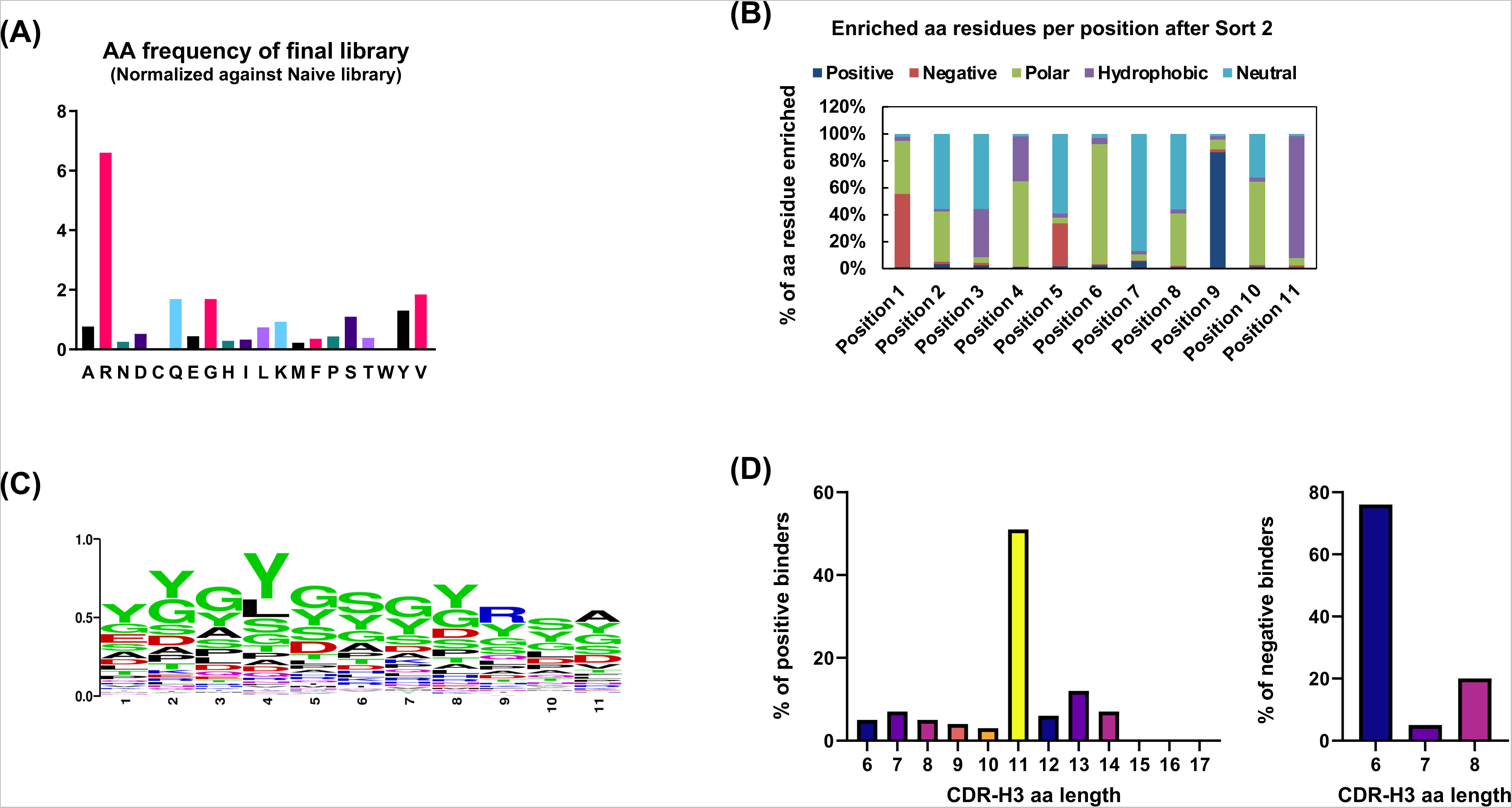
Next-generation sequencing of binders and non-binders to MMP-9. A) This bar graph displays the normalized frequency of amino acids in the final scFv library, compared to the naive library. Each bar’s height indicates the relative enrichment or depletion of a specific amino acid in the final library. Notably, certain amino acids, such as R, Q, G, Y, and V show significant enrichment, indicating selective pressure during the library evolution process. B) This bar graph represents the proportion of various amino acid types (Positive: R, K, H, Negative: D, E, Hydrophobic: A, I, L, M, F, W, V, Neutral: G, C, P, and Polar: N, Q, S, T, Y) at each position within the CDR-H3 region of the scFv variants with 11 aa lengths. Each bar corresponds to a specific position in the CDR-H3 region, showing the relative frequency of each amino acid type. The height of each colored segment within a bar indicates the prevalence of that particular amino acid type at the given position. This distribution helps in understanding the diversity and characteristics of the amino acids present in the CDR-H3 region, which can influence the binding affinity of the scFvs to the target MMP-9. C) The sequence logo illustrates the amino acid frequencies at each position within the CDR-H3 loop of the final scFv library with 11 aa residues. The height of each letter is proportional to its frequency at a given position, with taller letters indicating higher frequency. This visualization highlights the predominant amino acids at each position, revealing key residues that may contribute to the binding affinity. D) The distribution of CDR-H3 lengths among positive and negative binders reveals a notable trend: scFv binders often exhibit longer CDR-H3 regions, with 11 amino acid residues emerging as the predominant length within the final sorted library. Conversely, weak or non-binders typically possess shorter CDR-H3 sequences, with 6 amino acid residues being the most commonly observed length. This distinction underscores the importance of CDR-H3 length in determining binding affinity.

A deep analysis of all the unique sequences obtained for positive binders revealed that regardless of the length of CDR-H3, the combinations of Gly/Tyr/Ser are dominant (**Fig. 3C**). Overall, these results suggest that the enrichment of Tyr residues, which are larger in size, likely facilitates favorable contact with the MMP-9cd. Meanwhile, the presence of smaller Ser and Gly residues may contribute to suitable conformations beneficial to high-affinity binding. This observation also aligns with previous findings in the structures of Fabs selected from analogous minimalist libraries ^47,49^. Taken together, the results suggest that high-affinity binding is best mediated by Tyr in combination with small, flexible residues like Gly and Ser.

The NGS analysis was also employed to investigate the frequency distribution of CDR-H3 lengths among both scFv variants that were screened as positive binders and non-binders to MMP-9cd. Interestingly, CDR-H3 sequences with a length of 11 amino acids appeared the most frequently in the pool of positive binders. This was followed by CDR-H3 lengths of 13, 14, and 12 residues, respectively. In contrast, non-binders exhibited an enrichment of CDR-H3 lengths at 6 and 8 amino acids (**Fig. 3D**). These non-binding CDR-H3 sequences were also characterized by a high abundance of Leu, Phe, and Met (**Fig. S5**). These amino acids are classified as non-polar and uncharged side chains, which are typically considered less favorable for establishing strong molecular interactions with the target MMP-9cd. This observation suggests a potential correlation between CDR-H3 length and the binding affinity of isolated clones. The enrichment of shorter CDR-H3 lengths in non-binders, coupled with the dominance of non-polar amino acid residues, indicates a reduced capacity for forming key contacts with the MMP-9’s binding pockets.

### Performance of the DL model for predicting MMP-9cd binding of scFv variants

Stratified 10-fold validation was performed on the training sequences. The cross-validation and out-of-sample (test) metrics for the LSTM classifiers trained on features extracted from ESM-2 (650MB, 3B, and 15B) and AntiBERTy LPLMs (**Fig. 4**). The precision indicates the percentage of CDRH-3 sequences predicted by the model to be binders that were experimentally verified as binders. The recall indicates the percentage of experimentally verified binders the models detected. The F1 score is the harmonic mean of precision and recall. The LSTM model trained on features extracted from LPLMs is very effective at predicting the binding affinity of CDR-H3 to MMP-9 with precision close to or above 99%. The LSTM model trained on features extracted from the largest ESM model (ESM-2-15B) has the highest out-of-sample F1 score (99.11%). However, its F1 score exceeds that of the smallest model, AntiBERTy, with 26M parameters, by only 0.67%. Considering the memory usage and the speed of inference for larger models, one might prefer the smaller AntiBERTy model over the larger ESM models for this application.

**Figure 4.**
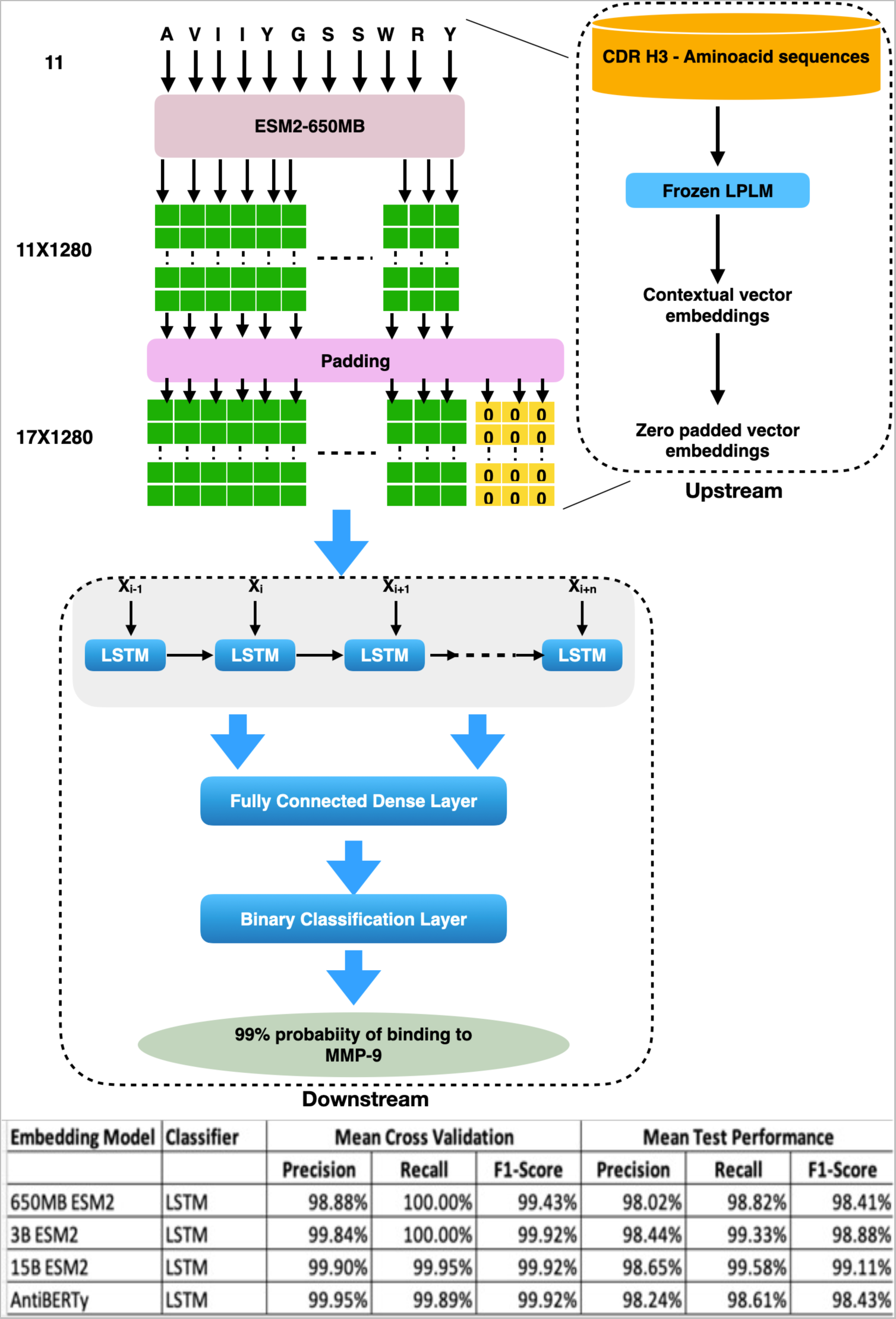
A computational model for predicting binding affinity of CDR-H3 to MMP-9. The pre-trained LPLMs are used to extract the vector representations (embeddings) for the CDR-H3s, which are padded to achieve a uniform length. These embeddings are used to train a downstream LSTM model with LSTM, dense, and binary classification layers. The REGA-3G12 CDR-H3 (AVIIYGSSWRY) length 11 is passed to the pre-trained LPLM ESM-2 650MB, which generates 11×1280 vector representations (embeddings) with 1280 vector representations for each amino acid. These embeddings are padded with zero vectors to achieve a uniform length of 17×1280, as 17 is the maximum CDR-H3 length in the samples provided to train the model. These zero-padded embeddings are passed to the downstream LSTM model, which is trained earlier to predict the binding affinity of CDR-H3 to MMP9. The model predicts a 99% binding probability to MMP-9cd for the REGA-3G12 CDR-H3 (AVIIYGSSWRY).

Force plots were created using the Shapley values generated by the DeepSHAP algorithm. The plot consists of arrows that represent each feature’s contribution, with their lengths proportional to the contribution’s magnitude. Positive contributions (shown in red) push the prediction towards binders, while negative contributions (shown in blue) push it towards non-binders. Indices were used to distinguish between the same amino acids in different positions of CDR-H3 (**Fig. 5A**). For instance, REGA-3G12 (CDR-H3: AVIIYGSSWRY) was predicted to be a positive binder, with the Gly6 and Val2 residues primarily contributing to this binding decision by the model. On the other hand, M0072 (CDR-H3: GAWYL), a non-binder scFv antibody to active MMP-9cd ^50^, was correctly predicted as a negative binder. The Leu5 residue mainly drove this non-binding prediction, as this residue frequently appeared in non-binder sequences discussed in the NGS analysis section. The residue-position mapping can also uncover specific interactions between residues and positions that are crucial for binding mechanisms to MMP-9cd (**Fig. 5B**).

**Figure 5.**
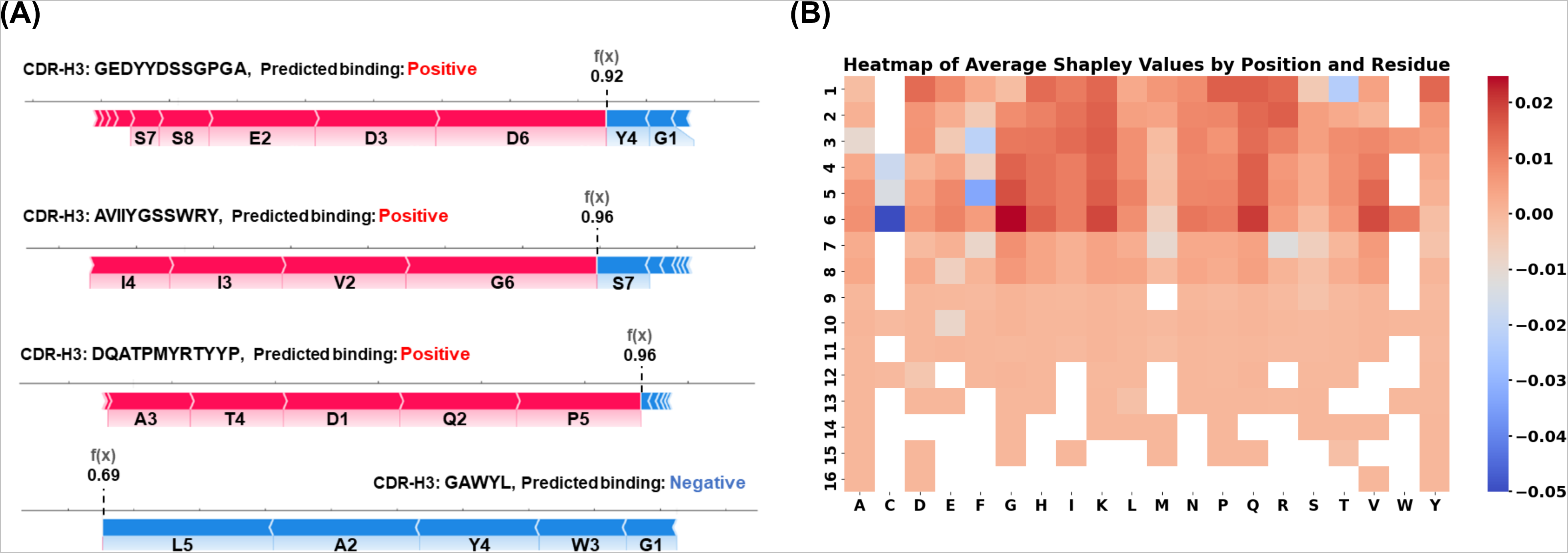
Residue-Position Mapping. A) The Shapley plot illustrates the final prediction of the machine learning model. Red arrows indicate specific amino acids that positively contribute to the binding prediction, while blue arrows represent amino acid residues that negatively participate in the binding prediction. B) The heatmap visually represents the impact of various amino acid residues at specific positions on binding affinity. Higher Shapley values (warmer colors) signify positive contributions to MMP-9 binding, highlighting crucial interactions between residues and positions. Conversely, lower Shapley values (cooler colors) indicate negative contributions. This can uncover specific interactions between residues and positions crucial for binding.

Analyzing Shapley values extracted from the ML model can be useful in identifying specific residues and positions in the CDR-H3 that enhance binding to MMP-9cd. The average (global) Shapley value for a specific position represents the mean contribution of that position to the scFv binding affinity to MMP-9cd, across different possible residues that could occupy that position (**Fig. S6A**). The positions with high positive average Shapley values (1 through 6) are critical and could be prime targets for mutations or engineering efforts aimed at improving binding affinity. Conversely, positions with low or negative Shapley values are associated with non-binding to MMP-9cd. The average (Global) Shapley value per amino acid residue represents the mean contribution of that residue to the scFv variants’ binding ability to the MMP-9cd (**Fig. S6B**). The bar chart reveals that the amino acid residues Cys, Phe, and Met, in general, have negative effects on binding irrespective of their locations. Positional variability captures the fluctuations in Shapley values for each position within the CDR-H3 region (**Fig. S6C**). This chart indicates that the contributions of positions 1 through 7 can vary greatly depending on the specific amino acid at these positions. This variability highlights the fact that these positions are likely crucial for binding interactions and could play a key role in the interaction mechanism.

### The screened scFv variants showed improvement in MMP-9cd binding

Following FACS screening to enrich scFv variants with improved binding affinity towards MMP-9 cd, individual clones from the final sorted library were isolated, grown, and induced for protein expression. Yeast cell surface display serves as a valuable indirect screening tool to assess two key properties of mutant proteins: enhanced thermal stability and efficient soluble secretion ^46,51^ shown by the expression level of scFv mutants displayed on the yeast. The scFv variant expression levels were monitored by labeling a c-Myc tag antibody conjugated to Alexa Fluor 488. A significant increase in expression levels (up to three-fold) and MMP-9cd binding up to six-fold was achieved compared to N-TIMP-1, a known tight binder of MMP-9cd ^37^(**Fig. 6A**, **Fig. S7**). This finding represents a significant achievement in developing high-affinity binders for MMP-9cd after two rounds of FACS. The conclusions can also be visually represented through dual scatter plots generated from flow cytometry data (**Fig. 6B**, **Fig. S8**). These plots demonstrate that the yeast population shifted towards higher MMP-9cd binding and expression values compared to N-TIMP1. This visual evidence further supports the significant increase in binding affinity and expression levels of the scFvs isolated from the synthetic scFv antibody library.

**Figure 6.**
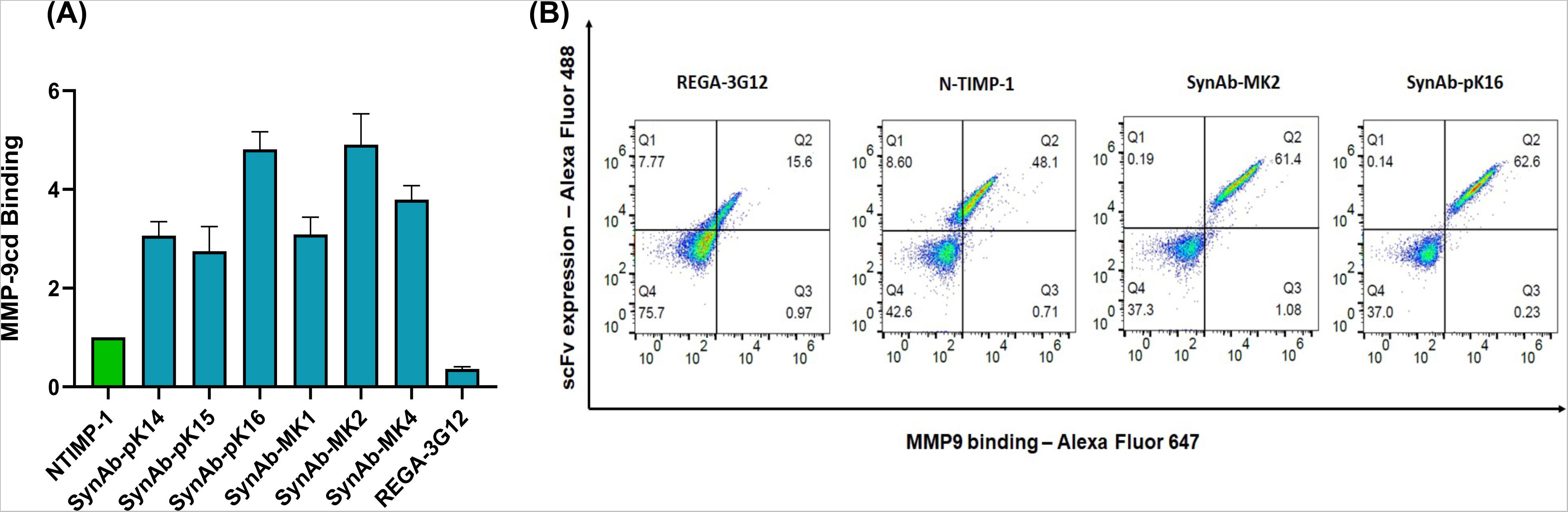
scFv variants isolated after two sequential FACS screening for MMP-9cd binding. A) Bar graph depicts the mean fluorescence intensity for His6x-MMP-9cd binding of scFv variants, adjusted for background and normalized to NTIMP-1 as a positive control for MMP-9cd binding. Yeast-displayed scFv variants were incubated with 300 nM soluble MMP-9cd protein in all experiments. Each data point represents the mean of triplicate samples, with error bars indicating the standard error of the mean (SEM). B) Flow cytometry scatter plots for several isolated yeast-displayed scFv variants exhibiting improved MMP-9cd binding activity are presented, with NTIMP-1 as a reference. The x-axis (APC channel) represents binding to His6x-MMP-9cd (300 nM), while the y-axis (FITC channel) indicates scFv expression levels.

After the Sanger sequencing of isolated clones (**Table. 1**), it can be observed that clones containing negatively charged residues such as Asp, and positively charged residues like Lys, His, and Arg exhibit higher expression compared to those that do not contain these residues. Overall, an increase in charged and hydrophilic residues, along with a decrease in hydrophobic residues, improved solubility of scFv variants consistent with previous observations in this area^52,53^.

**Table 1.**
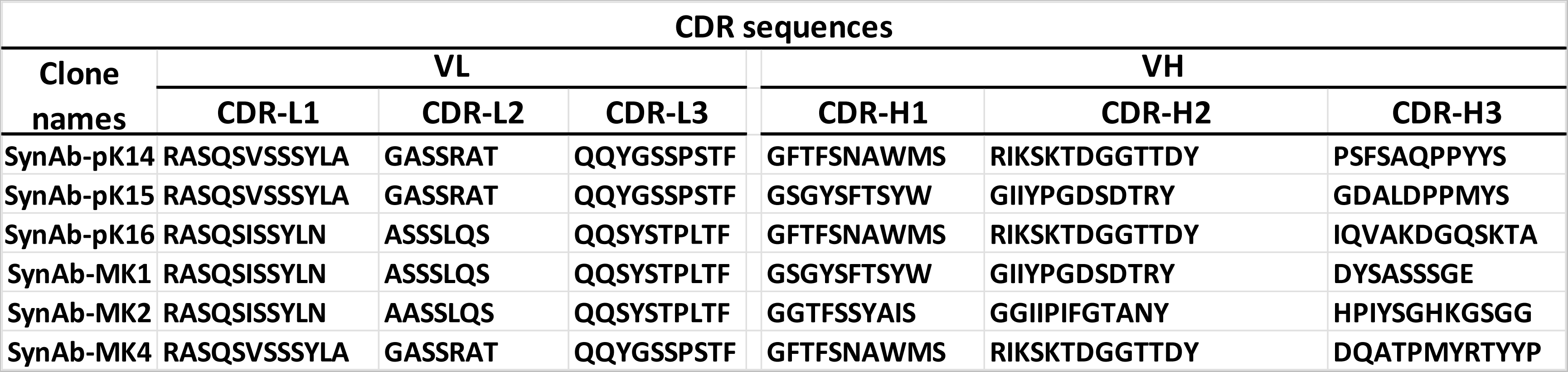
Sanger sequencing of isolated scFv antibodies. Sequences of the different CDR regions in the light and heavy chains are shown in the table. Kabat numbering is used for numbering the residues in each antibody.

### Structural studies of scFv variants with improved MMP-9 binding shows CDR-H3 contacts with MMP-9cd active site

AlphaFold2 was used for the structural modeling of protein complexes of all engineered scFvs against MMP-9cd. AlphaFold2 is a groundbreaking tool for predicting the structures of protein complexes. It utilizes a complex deep learning architecture, including the Evoformer module and Invariant Point Attention (IPA), specifically designed to handle the sophisticated interactions between proteins ^54^ Unlike traditional methods that analyze individual chains separately, AlphaFold2 Multimer excels by integrating all interacting chains within a single prediction run, effectively simulating the entire complex. This holistic approach leads to superior accuracy in predicting three-dimensional structures ^55^. Furthermore, AlphaFold2 Multimer leverages information from existing protein structures deposited in the Protein Data Bank (PDB) to enhance its predictions. Additionally, it refines the generated models using well-established molecular dynamics tools like Amber. More importantly, AlphaFold2 Multimer provides users with confidence metrics, such as the predicted Local Distance Difference Test (pLDDT), predicted template modeling (pTM), and interface predicted template modeling (ipTM) to assess the reliability of the predicted interactions. This combination of features makes AlphaFold2 Multimer an exceptional tool for predicting protein complexes, particularly valuable for understanding complex interactions like those between antibodies and antigens ^56^. However, the need for co-evolution data restricts the accuracy of predicting antibody-antigen interactions to some extent since antibodies bind strongly to antigens not due to co-evolution but owing to processes such as somatic hypermutation and affinity maturation ^57^.

Analysis of confidence metrics for the modeled isolated scFvs revealed high confidence in the accuracy of the model structures. For instance, all structures have a pLDDT score exceeding 90, indicating AlphaFold’s high confidence in the positions of each atom within the predicted structures. Furthermore, the pTM score, which assesses both the accuracy of the entire predicted protein structure and its similarity to a known reference structure, is above 0.85 for all structures, signifying a high degree of accuracy.

Following the structural modeling of MMP-9cd and the engineered scFv variants protein complexes, the protein surface topology and CDR-H3 charge were also investigated (**Fig. 7**). MMP-9cd’s active site possesses a characteristic negative charge and a concave geometry. Notably, compared to other MMPs, the active site of MMP-9cd exhibits a flatter conformation.

**Figure 7.**
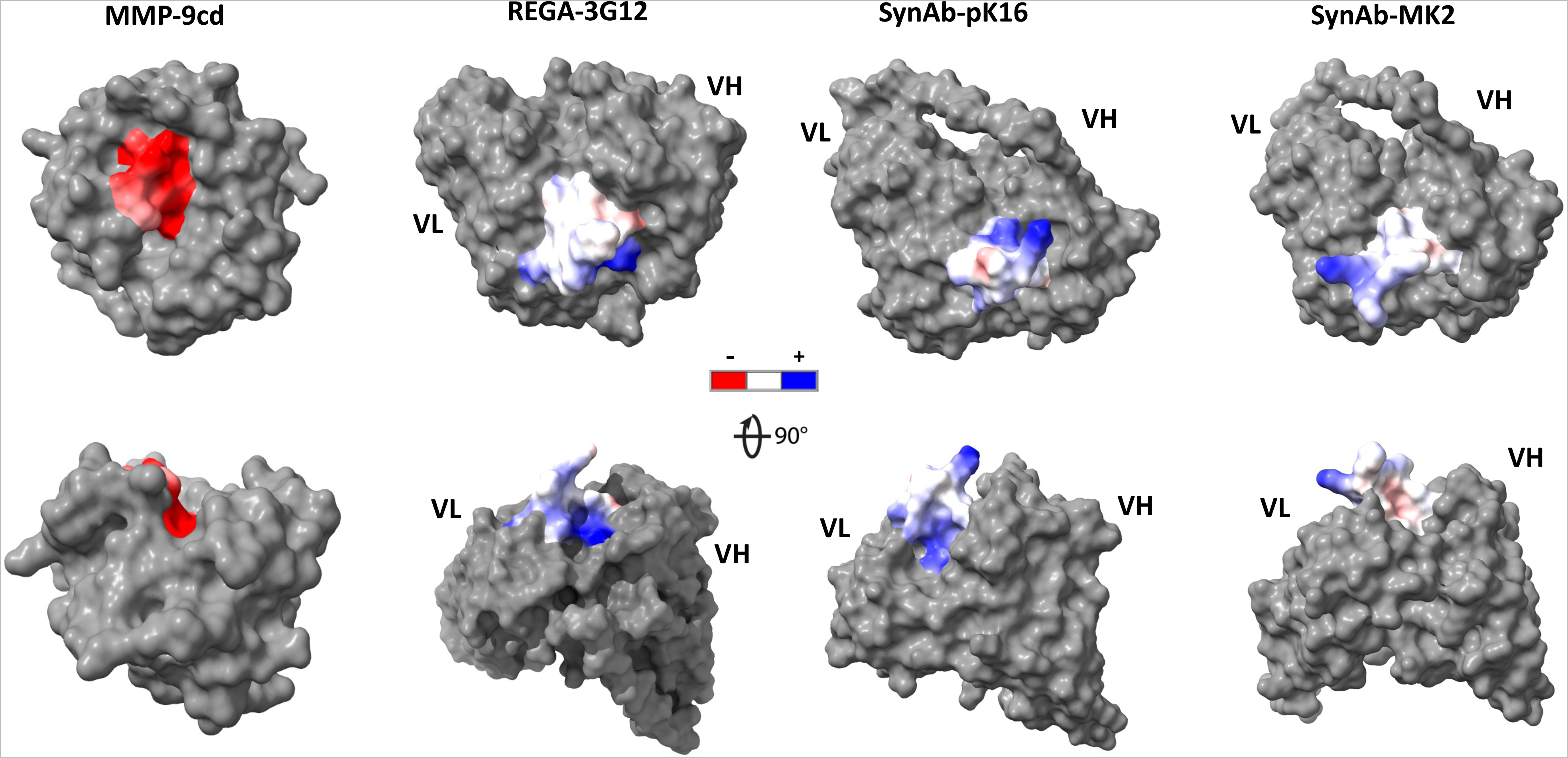
Surface charge distribution of CDR-H3 on different scFvs and MMP-9cd. The surface charge is indicated by color, where red represents negatively charged regions, blue positively charged regions, and white neutral regions. The 90° rotation provides a comprehensive view of the charge distribution, highlighting the three-dimensional electrostatic landscape and geometry of each protein’s CDR-H3. The negatively charged area in the active site of MMP-9cd is prominent, suggesting a strong electrostatic interaction potential with positively charged CDR-H3 sequences. As a positive control, REGA-3G12’s CDR-H3 domain has a neutral to positive charge distribution suited for electrostatic interactions with MMP-9cd. Its protruding geometry complements MMP-9’s active site, potentially optimizing binding. In this way, both isolated top clones SynAb-pK16 and SynAb-MK2 show similar CDR-H3 electrostatic charge distributions and geometries, suggesting specialized adaptations for binding to MMP-9cd.

This distinct feature presents a favorable opportunity for the directed engineering of scFv variants, particularly focusing on optimizing CDR-H3 length and amino acid compositions of murine or human antibodies for enhanced binding to this specific region. REGA-3G12 is considered a selective inhibitor of MMP-9cd, featuring 11 amino acid residues in the CDR-H3 region ^22,23^. As a positive binder and inhibitor of MMP-9, the REGA-3G12’s CR-H3 loop has a positive surface charge, which explains its favorable binding affinity to the negatively charged region of the active site and neighboring regions. Additionally, the convex shape of CDR-H3 in REGA-3G12 is a proper fit to the concave shape of the MMP-9cd active site, allowing it to fit in in the active site effectively. The same positive charge and the convex shape of CDR-H3 were observed with most of the engineered scFv variants, especially SynAb-pK16, and SynAb-MK2, which showed up to 5-fold higher binding affinity compared to N-TIMP-1. They have 12 amino acid residues in the CDR-H3 regions and a positively charged surface. The length and amino acid composition of this region also confer a convex shape to these isolated scFvs, making them promising candidates for targeting MMP-9cd. These observations help explain why most of the frequently enriched amino acid residues are charged and polar, and why the length of CDR-H3 tends to be longer in strong binders compared to non-binders or weak binders, which have shorter CDR-H3 regions with more non-polar residues. This pattern shows that the charge in the variable regions of antibodies, especially in the CDR regions, affects both their ability to bind to antigens and their overall stability and folding. Charged regions contribute significantly to these properties, with mutations that introduce or remove charges impacting folding efficiency, stability, and binding affinity under different conditions ^58^.

On the other hand, by examining the interaction of only the CDR-H3 regions of isolated scFvs with MMP-9cd, specific positions and residues responsible for driving MMP-9cd binding, and possibly the direction of binding towards the active site, can be identified. Protein complexes predicted by AlphaFold multimer for REGA-3G12, a known positive binder, and SynAb-pK16 and SynAb-MK2, the two highest binders to MMP-9, reveal interesting interactions between the scFvs and MMP-9cd (**Fig. 8**). As indicated by the analysis of general Shapley values, the first six positions of CDR-H3 are shown to contribute significantly to positive binding, with a particular emphasis on positions 6,5, and 1. Interestingly, for all these scFvs, position 6 is found to play a key role in binding to the active site residues or neighboring residues of MMP-9cd, specifically involving Gly104, Asp233, and Gly231 in REGA-3G12, SynAb-pK16, and SynAb-MK2, respectively. Regardless of position number 6, it can be observed that positions number 1, 4, and 5 also play roles in making interactions with MMP-9cd in these scFv variants, highlighting the significance of these positions in directing the binding towards MMP-9cd. In summary, the findings emphasize the role of CDR-H3, particularly the first six positions, in driving the binding between scFvs and MMP-9cd. It highlights a potential hot spot residue at position 6 for interaction with MMP-9cd and underlines the importance of other residues (positions 1, 4, and 5) for proper binding orientation.

**Figure 8.**
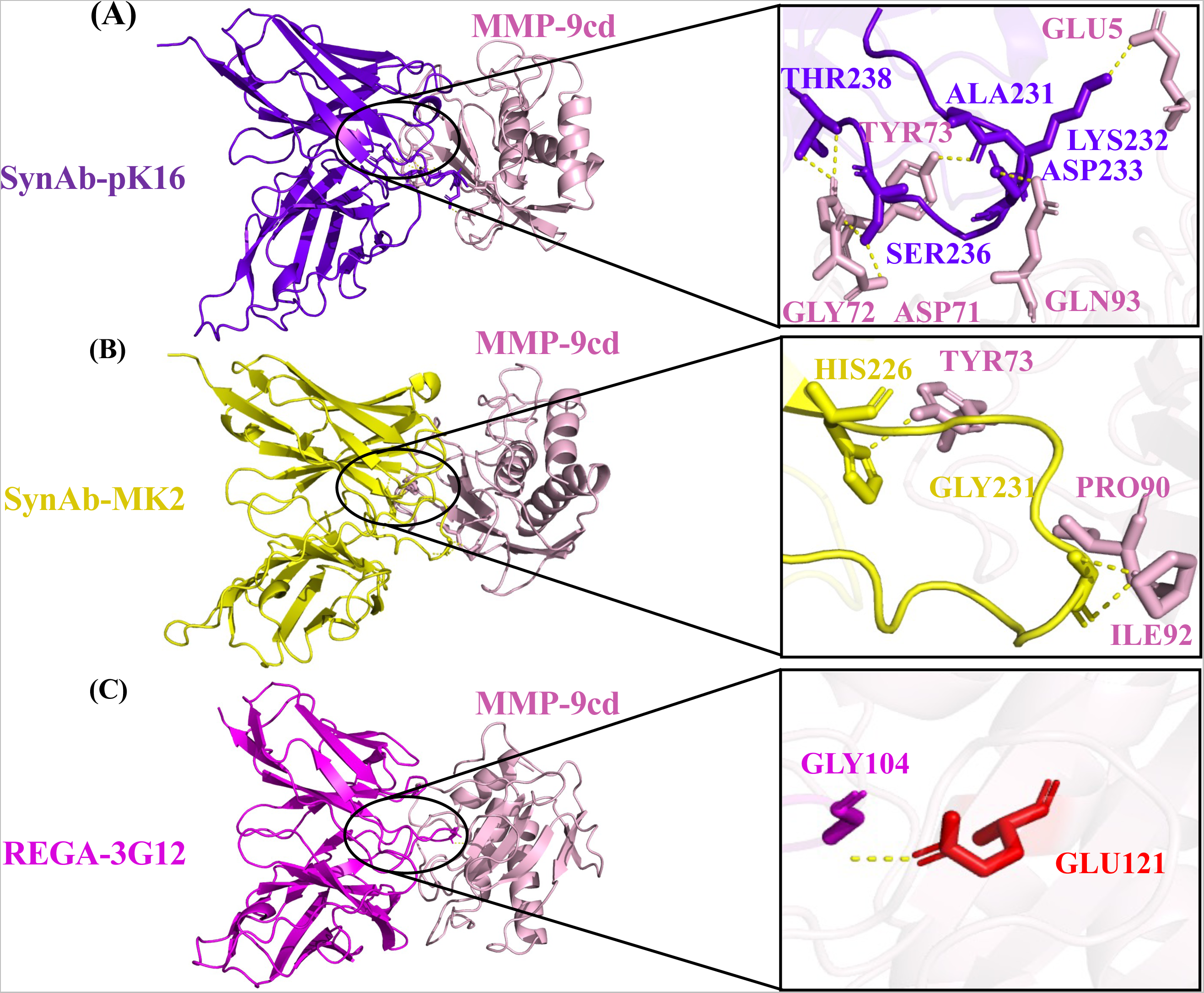
Structural analysis of scFv clones bound to MMP-9cd. Highlighting the role of CDR-H3 residues in binding to MMP-9cd from AlphaFold2 protein complexes. A) The CDR-H3 residues of SynAb-pK16 (highlighted in purple) show binding interactions with MMP-9cd (highlighted in light pink). Specifically, Ala231, Lys232, Asp233, Ser236, and Thr238 are involved in these interactions. B) SynAb-MK2 (highlighted in yellow) binds to MMP-9cd (highlighted in light pink) through its CDR-H3 residues, specifically via Gly231 and His226. C) REGA-3G12 (highlighted in dark pink) binds to MMP-9cd (highlighted in light pink) through the Gly104 residue in its CDR-H3.

## Discussion

Directed evolution using yeast surface display of a synthetic antibody library previously engineered for nonspecific binding ^24^was used to screen, isolate, and analyze the scFv antibody variants with improved binding to MMP-9cd compared to endogenous MMP-9 inhibitor, N-TIMP-1, and other MMP-9 antibodies such as REGA-3G12 ^22,23^.

The role of CDR-H3 in binding to MMP-9 was investigated using experimental and computational models, considering CHR-H3’s significance in antibody-antigen interactions due to both length and amino acid composition. The next-generation DNA sequencing analysis found patterns in the CDR-H3 of scFvs binders with polar residues such as Ser, and Tyr being dominant in frequency. The most favorable length of CDR-H3 for MMP-9 binding was found to be 11 amino acids which was confirmed to be the most effective in binding to the MMP-9 catalytic site using computational modeling and structural analysis. Further, importance of CDRH-3 charge was highlighted as the selected scFv variants showed positive surface charges which match the negative charge in the MMP-9cd active site.

A machine learning strategy based on recent advances in developing protein language models to predict protein structure and function was employed to develop a model that predicts scFv variants binding to MMP-9 based on CDR-H3 sequences, recognizing it as the key driver of binding. The developed deep learning language model was used to predict the binding of other MMP-9 antibody binders and non-binders which sequences were not used in fine-tuning the models with high accuracy proving its potency to be used as a predictor tool for designing new antibodies for binding to protein targets such as MMP-9.

The MMP activity could also be blocked by targeting the neighboring regions outside of the catalytic domain, known as exosites ^59^. Unlike the catalytic domains, which are similar across MMPs, different MMPs have distinct exosites. Targeting exosites using synthetic antibodies was previously used to provide selective inhibition of specific MMPs which led to the development of highly selective mAbs targeting MMP-9, such as AB0041 and AB0046, as well as their humanized version, GS-5745 ^60^. Using the developed language model tools to optimize key regions of scFv antibodies focusing on CDR-H3 regions could facilitate overcoming the limitations in targeting specific MMPs.

## ABBREVIATIONS

MMP: matrix metalloproteinase
FACS: fluorescent-activated cell sorting
scFv: single-chain antibody fragment
TIMP: tissue inhibitors of metalloproteinases
NLP: natural language processing
LPLM: large protein language model
LSTM: long-short-term memory
SHAP: SHapley Additive exPlanations
ML: machine learning
ESM: evolutionary scale modeling
PCA: principal component analysis
t-SNE: t-distributed Stochastic neighbor embedding
MLM: Masked Language Modeling.

## Acknowledgment

Figures 1 and 2A were created with BioRender.com. We would like to thank Prof. Dane Wittrup (Chemical Engineering-MIT) for the generous gift of the nonspecific synthetic antibody library. M. R.-S. had funding support from NIH R03AG070511 and NIH R21HD109743.

## Author contributions

M.R.-S., E. K.-B; conception, M.K.; performed the experiments, I.K.: performed machine learning models, S. K., M.K.: performed computational analysis, M.K., I.K., E. K.-B, M.R.-S., data analysis, writing, and editing the manuscript drafts. M.R.-S. funding and supervision.

## Supplement Figures

**Fig S1.**
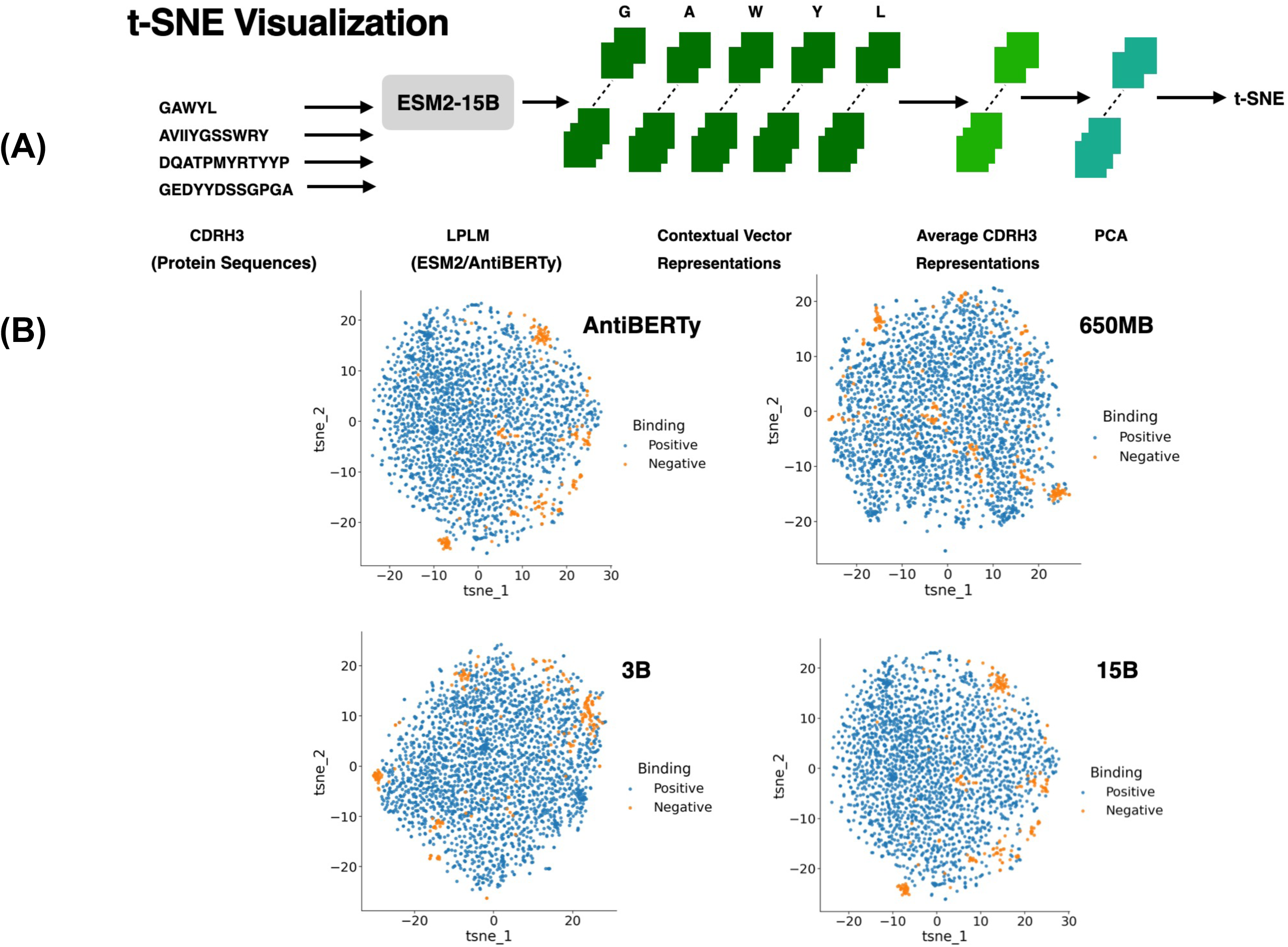

**Fig S2.**
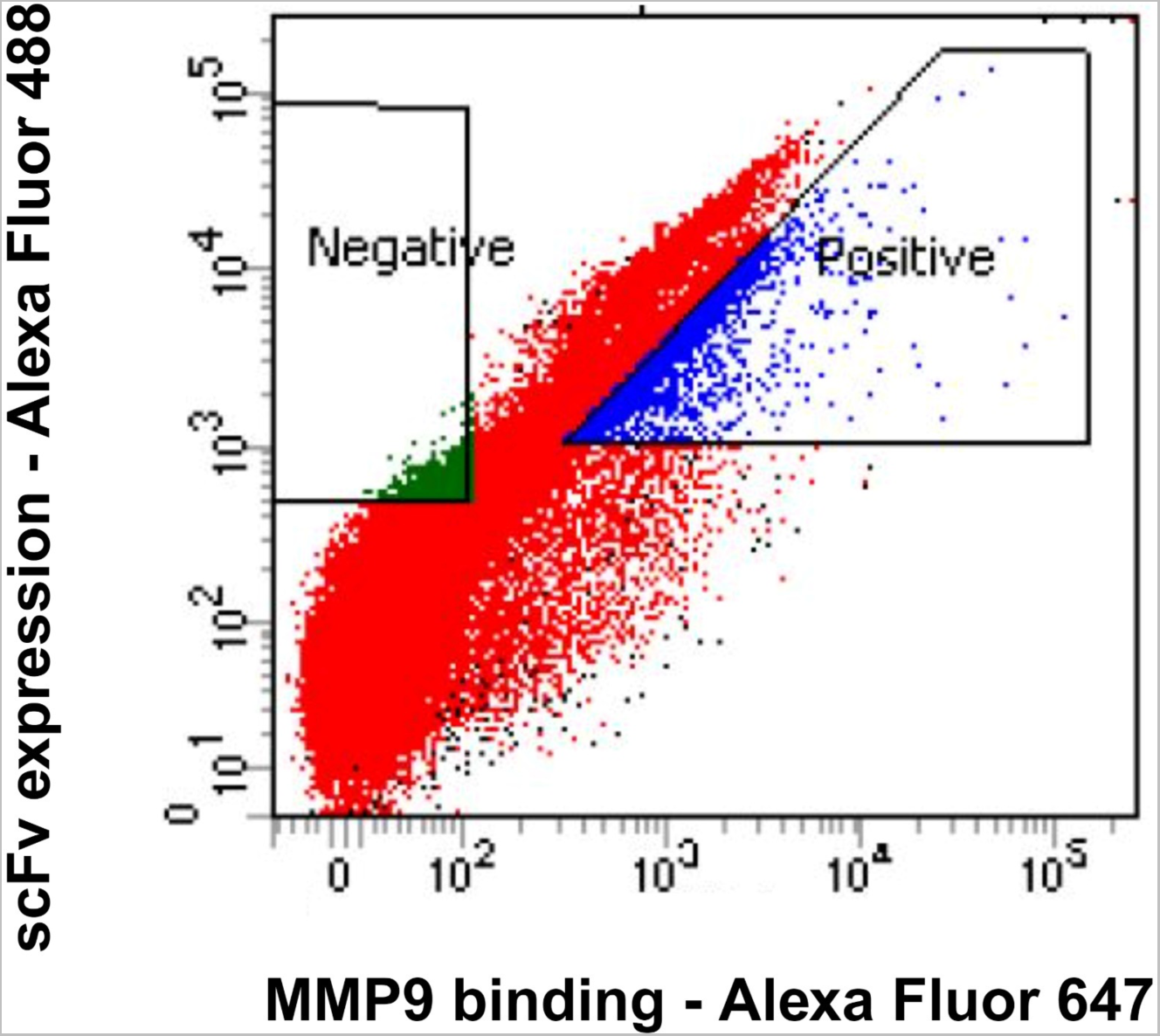

**Fig S3.**
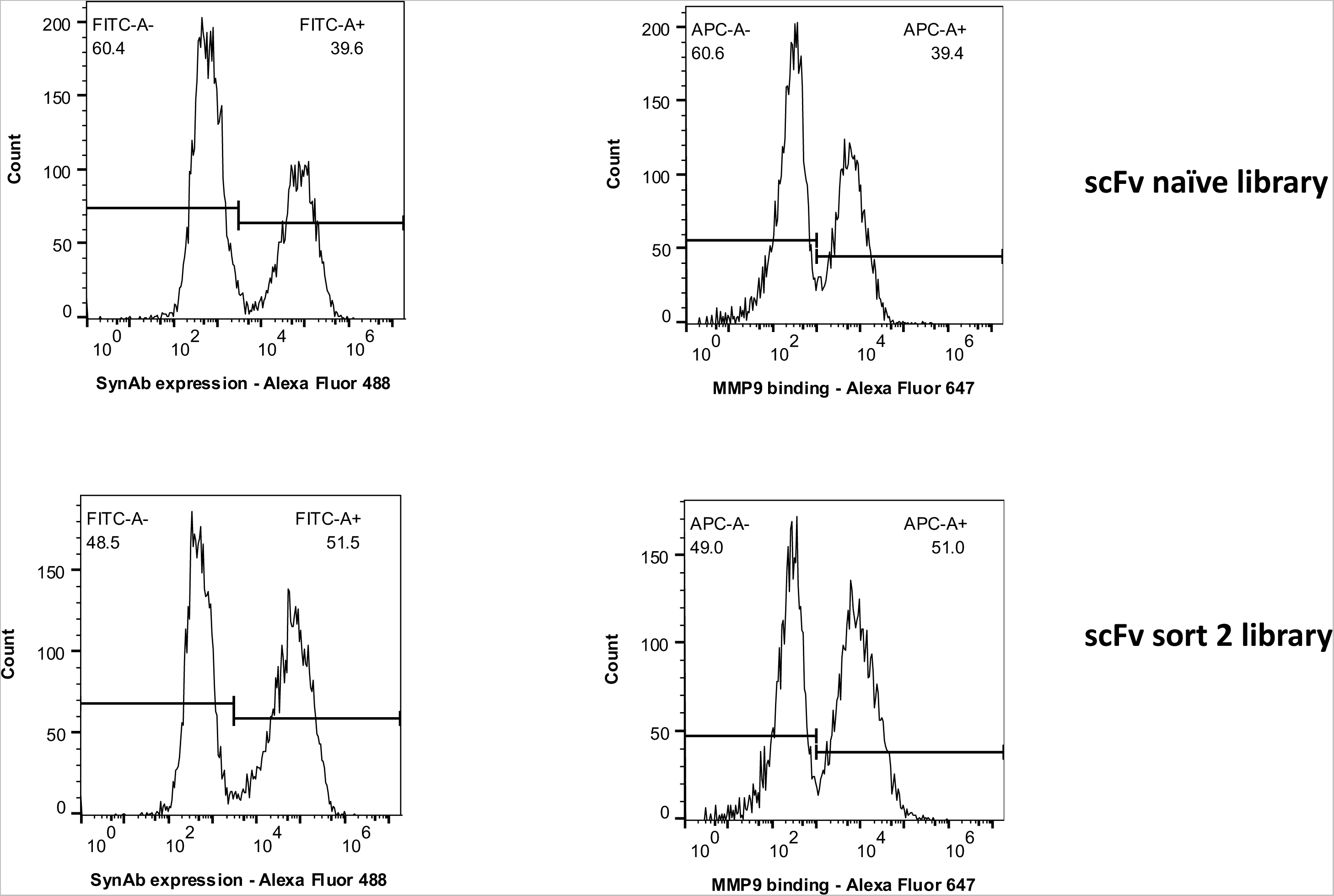

**Fig S4.**
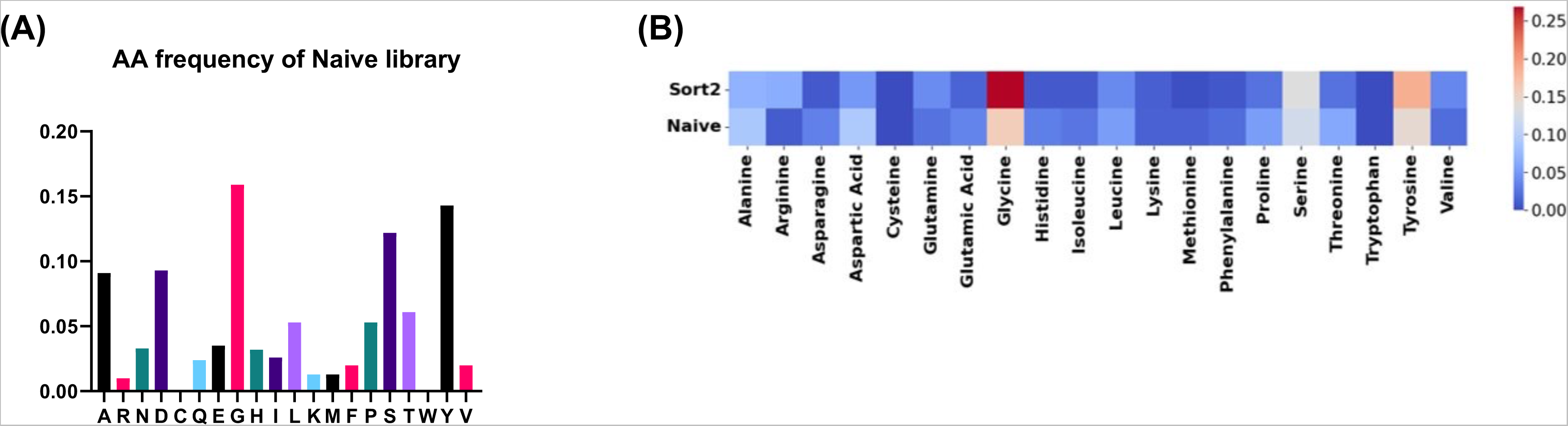

**Fig S5.**
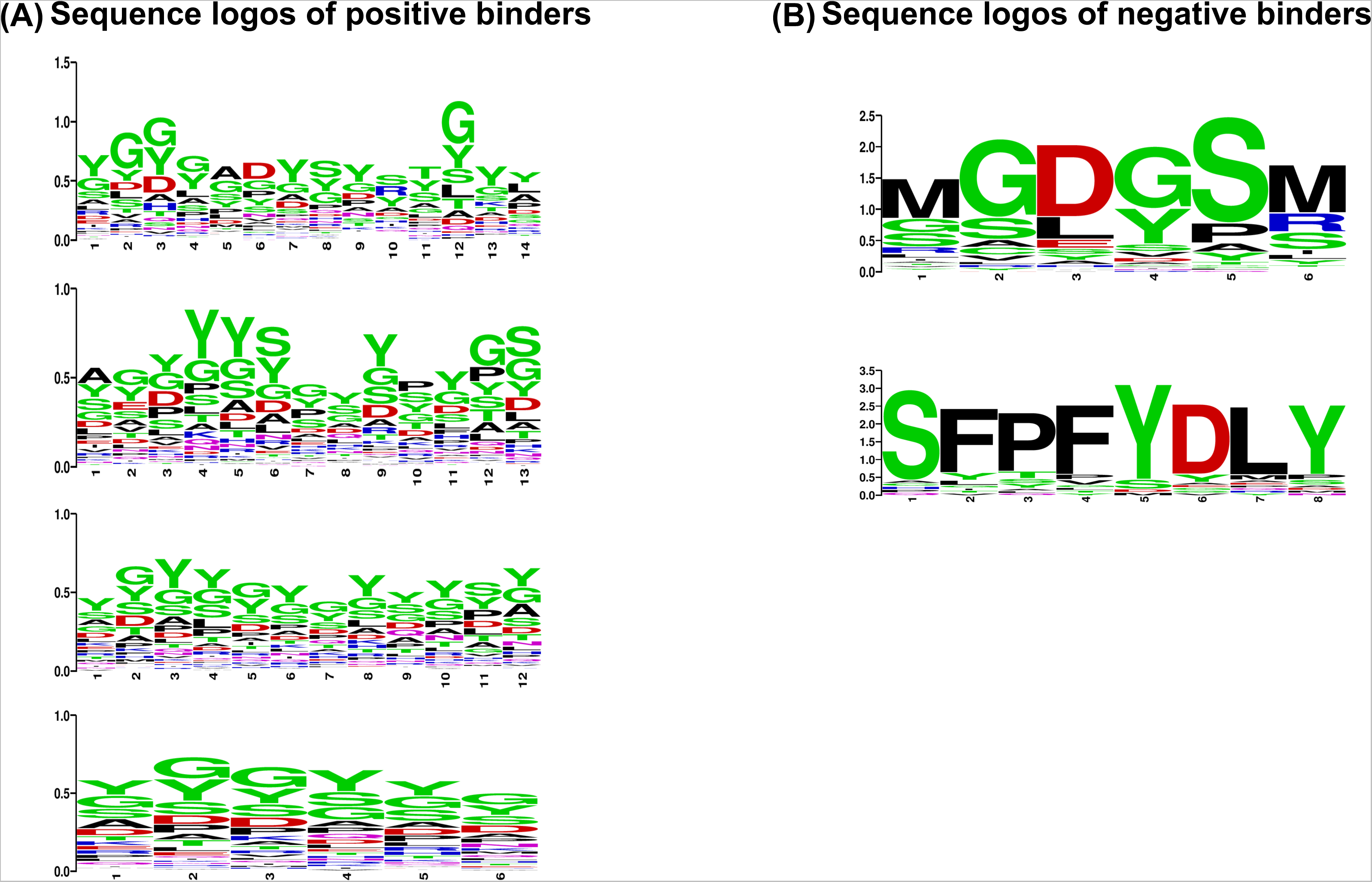

**Fig S6.**
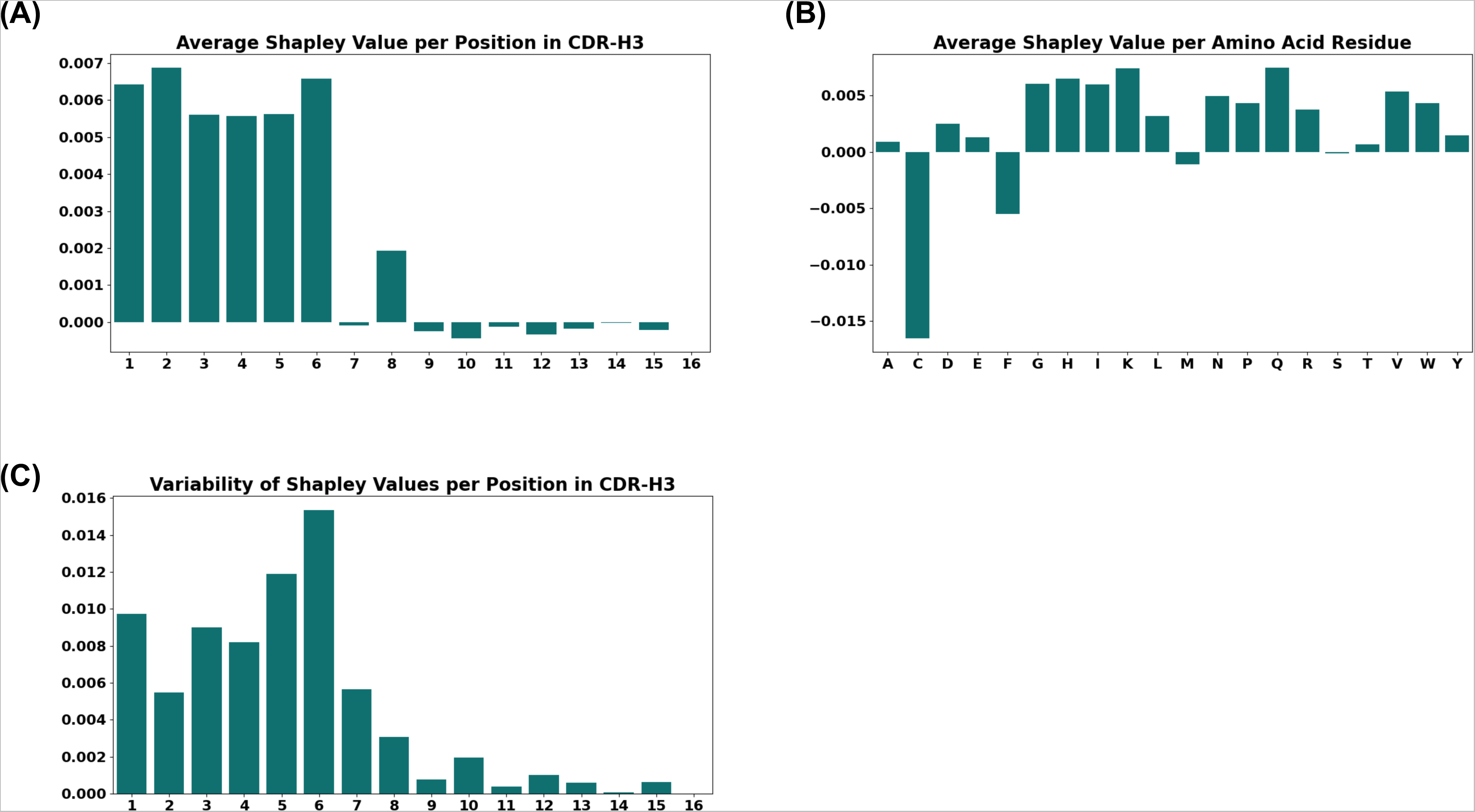

**Fig S7.**
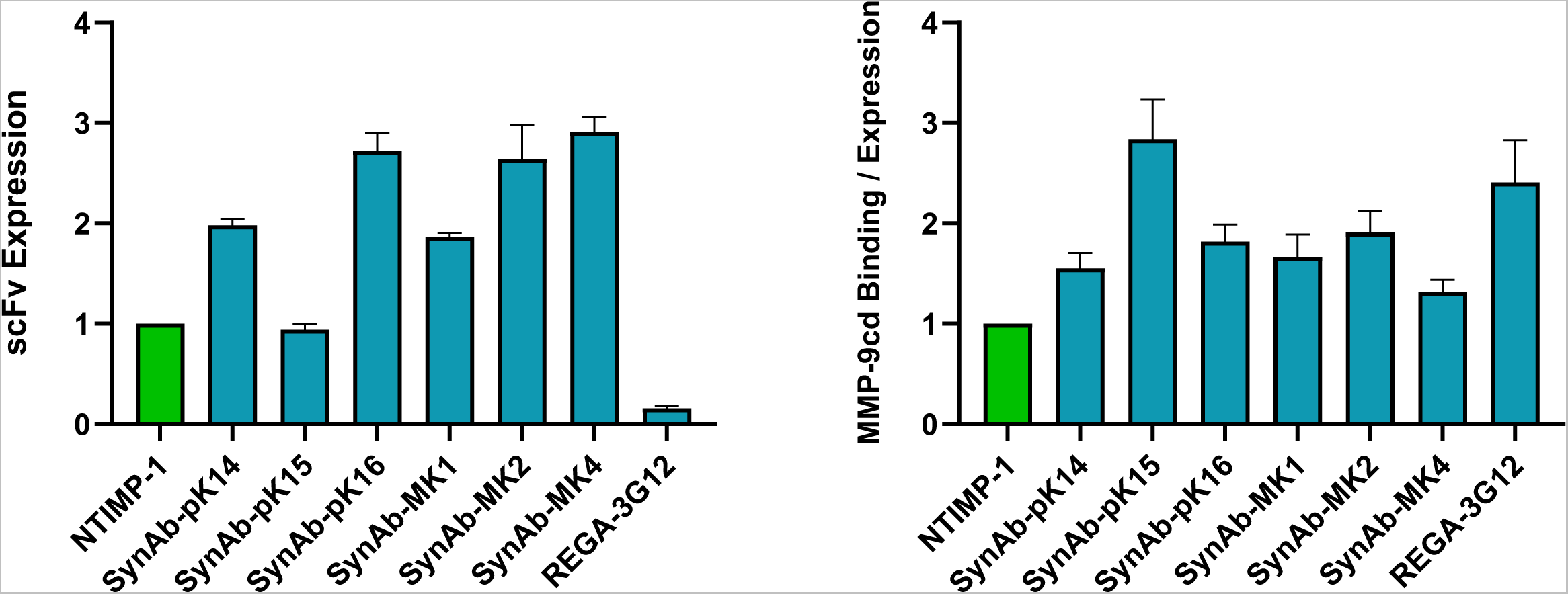

**Fig S8.**
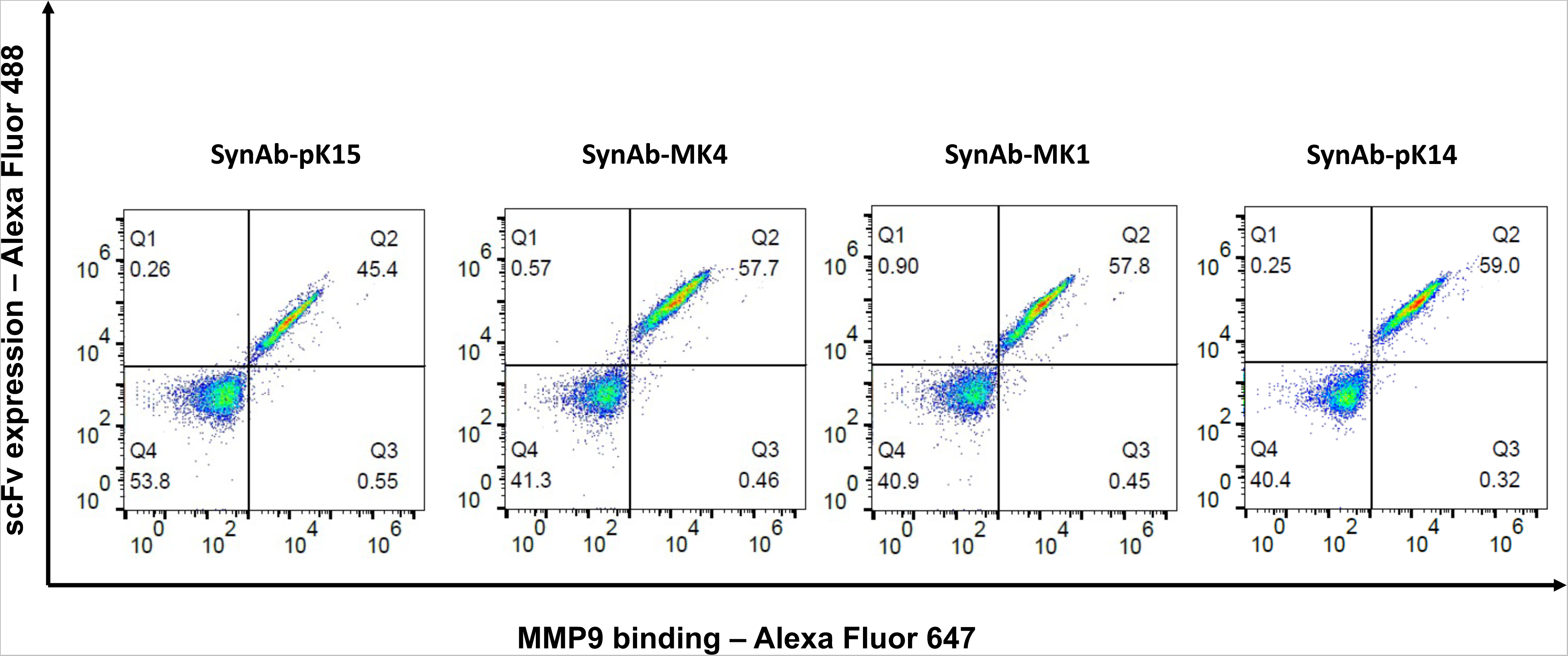

