## Supplemental Figure Legends for "Elucidating key determinants of engineered scFv antibody in MMP-9 binding using high throughput screening and machine learning"

**Figure S1. Low dimensional visualization of features extracted for CDR-H3 sequences using ESM-2 and AntiBERTy LPLMs.** AA sequences of CDR-H3s are passed to pre-trained large protein language models like ESM2-15B. The ESM2-15B creates 5120 representations for each amino acid in the CDR-H3. Then, an average is taken over the entire CDR-H3 to get a single 5120 representation for a given CDR-H3, which is compressed using the Principal Component Analysis (PCA) technique to get t-SNE visualization.

**Figure S2. Cell sorting for binders and non-binders using FACS.** Positive and negative gates are employed to separate binders and non-binders among a population of scFv variants. A diagonal gate is used to collect binders targeting MMP-9cd, identifying variants with high signal values for both expression (Alexa Fluor 488) and binding (Alexa Fluor 647). Conversely, a rectangular gate is employed to gather non-binders that exhibit expression but possess low binding signal values.

**Figure S3. Comparison of expression and binding between Naïve and Sort2 scFv libraries.** The figure displays flow cytometry histograms representing scFv expression and binding levels. Two separate graphs are shown: one for scFv expression (labeled "FITC-A") and the other for MMP-9cd binding (labeled "APC-A"). Each histogram has two distinct peaks, corresponding to positive and negative cell populations. Thresholds are marked to identify cells above or below specific expression or binding levels compared to unlabeled yeast cells as negative control.

**Figure S4. Enrichment of amino acid residues in CDR-H3 loop**. A) Frequencies of individual amino acids in CDR H3 loops of Naïve scFv library. B) The enrichment ratio of diversified residues, as determined through FACS, is visually represented by a heat map. This enrichment ratio is computed based on NGS read counts corresponding to each CDR-H3 loop. Changes in the frequency of amino acid residues are depicted through color variations on the heat map, with warmer colors signifying an enrichment of specific residues and cooler colors indicating their depletion after two rounds of FACS.

**Figure S5. CDR-H3 Sequence Logo of Key Residues for MMP-9cd Interaction.** The sequence logo illustrates the amino acid frequencies at each position within the CDR-H3 loop of the final scFv library, reflecting the most frequent amino acid length for both positive and negative gates. The height of each letter corresponds to its frequency at a specific position, with taller letters representing higher frequencies. This visualization highlights the predominant amino acids at each position, revealing key residues that may contribute to the interaction with MMP-9cd.

**Figure S6. Shapley value analysis.** A) The average Shapley value per position represents the mean contribution of that position to the binding affinity of the scFv to MMP-9 across different residues. Positions with high average Shapley values are crucial for binding affinity while low or negative values do not contribute significantly to the binding process. B) Average Shapley values per residue refer to which residues generally have positive or negative impacts on binding affinity, without considering where they are located within the sequence. It provides insights into which amino acids might be preferred or avoided in the scFv design. C) Positional variability refers to the extent of fluctuation in Shapley values for each position within the CDRH3 region. High variability at a position indicates that its contribution to binding affinity can significantly differ depending on the amino acid present. Such variability suggests these positions might be critical for binding interactions, potentially serving as key determinants in the binding mechanism.

**Figure S7. scFv variants isolated after two sequential FACS screenings for expression levels and MMP-9 binding per expressed variants on yeast cells.** Bar graphs illustrate the average fluorescence intensity for c-myc expression, His6x-MMP-9cd binding/expression ratio of scFv variants. The data have been adjusted for background and normalized to NTIMP-1, serving as a positive control for MMP-9cd binding. In all experiments, yeast-displayed scFv variants were incubated with 300 nM of soluble MMP-9cd protein. Each data point represents the mean from triplicate samples, with error bars representing the standard error of the mean (SEM).

**Figure S8.** Flow cytometry scatter plots are shown for isolated yeast-displayed scFv variants that demonstrate enhanced binding activity to MMP-9cd, with NTIMP-1 included as a reference. The x-axis (APC channel) displays binding to His6x-MMP-9cd at 300 nM concentration, while the y-axis (FITC channel) indicates the expression levels of scFv.
